# Strain-specific thermotolerance, UV-C tolerance, and biofilm formation on clinically relevant plastic substrates in the emerging opportunistic pathogen *Rhodotorula mucilaginosa*

**DOI:** 10.64898/2026.08.19.745829

**Authors:** Yifei Chen, Isabel A. Jimenez, Arturo Casadevall, Piotr R. Stempinski

## Abstract

*Rhodotorula mucilaginosa* is an emerging opportunistic fungal pathogen increasingly associated with catheter-related bloodstream infections. Although biofilm formation is considered a major virulence trait for *R. mucilaginosa*, factors contributing to biofilm persistence on medical devices remain poorly understood. Here, we characterized the thermotolerance, biofilm formation, UV resistance, and cell surface hydrophobicity profiles of eight *R. mucilaginosa* strains representing clinical and non-clinical (laboratory, environmental, and marine mammal) isolates. All strains grew optimally at 30°C and exhibited restricted growth at 35°C and 37°C, although one environmental isolate maintained robust growth at 37°C. All strains exhibited moderate to high cell surface hydrophobicity. We then assessed biofilm formation for each strain, including adherence to two different plastic substrates, development of biofilm biomass, comparison of biofilm metabolic activity, and the effects of temperature on biofilm formation. Under static conditions, biofilm biomass of most isolates on 96-well polystyrene plates was greatest at 24°C. Clinical isolates generally maintained higher biofilm metabolic activity at 37°C than nonclinical isolates, while at lower temperatures, clinical and non-clinical isolates did not differ significantly in metabolic activity. All strains readily formed biofilms on polyurethane intravenous catheters under dynamic conditions, as confirmed by scanning electron microscopy and metabolic activity.

While planktonic cells already displayed substantial UV-C tolerance, biofilm-associated cells remained viable following exposure to UV-C doses up to eightfold higher than those that impaired planktonic growth. These findings document differences in thermotolerance and biofilm formation by isolate origin and identify biofilm formation as a major factor promoting persistence of *R. mucilaginosa* on clinically relevant materials and reduced susceptibility to UV-C sterilization.

## Introduction

*Rhodotorula* is a basidiomycetous environmental yeast found in various ecological niches, including soil, water, plants, animals, and food products (1, 2). *R. mucilaginosa* is frequently encountered in healthcare settings where it colonizes and forms biofilms on indwelling medical devices. It is globally ubiquitous and can be found in remote environments, including submarines and the International Space Station (3, 4). Various species of *Rhodotorula* can be identified as a part of the natural microbiota of human skin (2, 5). In recent years, increasing prevalence of *Rhodotorula* infections has resulted in classification of this species as an emerging fungal pathogen with the ability to colonize and cause fungemia in susceptible hosts (6–9). The vast majority of reported *Rhodotorula* clinical infections are caused by *R. mucilaginosa*, *R. minuta,* and *R. glutinis* (9–13). Most of these infections are associated with colonization of medical or hygienic devices, especially central and peripheral venous catheters, which provide a favorable surface for fungal attachment and biofilm formation (9, 13–16). After establishing stable surface adhesion, catheter-associated biofilms can promote persistent infection and increase the risk of septic fungemia (17–19).

Fungal biofilms are complex structured communities of cells covered by an extracellular matrix that can attach to various biotic or abiotic surfaces (20, 21). The main components of biofilms are cells, extracellular polysaccharides (EPS), extracellular DNA, proteins, and lipids (17, 22). Microbial biofilms can act as a protective barrier against antimicrobial agents and environmental stressors, including reactive oxygen species (ROS) and Ultraviolet light (UV) radiation, enhancing the survival and persistence of fungal colonies (23, 24). Biofilm formation on indwelling medical devices, including intravenous catheters, urinary catheters, and prosthetic implants, poses a major clinical challenge by promoting long-term fungal presence and serving as a reservoir for persistent or recurrent fungal infections (16, 20, 21). Management of biofilm-associated infections on medical devices requires removal of the contaminated material, which can negatively impact treatment and increase costs associated with therapy (20, 25–28).

In this study, we evaluated adaptation, growth, and biofilm formation at various clinically relevant temperatures of eight *R. mucilaginosa* strains, including a laboratory reference strain, three environmental strains, and three clinical isolates. We also assessed the UV-C tolerance of static fungal biofilms in these strains. Finally, we evaluated the biofilm formation of selected *R. mucilaginosa* strains on the surface of polyurethane catheters under dynamic conditions to simulate the clinical environment of an indwelling intravenous catheter.

## Results

### Thermotolerance in *Rhodotorula mucilaginosa* strains

*R. mucilaginosa* strains used in this study originated from diverse ecological and clinical sources, including a reference laboratory strain (ATCC 9449), an environmental strain isolated from a mosquito, *A. funestus* (Af401), environmental strains isolated from nasal swabs collected from harbor seals (MM20 and MM127), and strains of clinical origin (0438, 5429, 1364, and 9562). To evaluate their thermotolerance and adaptation to growth at elevated temperatures, strains were assessed by spot dilution assay on YPD agar plates at 24°C, 30°C, 35°C, and 37°C for 48 h (Fig. 1A). These temperatures were selected to represent ambient environmental conditions (24°C), the optimal growth temperature for the yeast (30°C), peripheral host temperatures such as those encountered on the surface of the skin (35°C), and core human body temperature (37°C).

**Figure 1.**
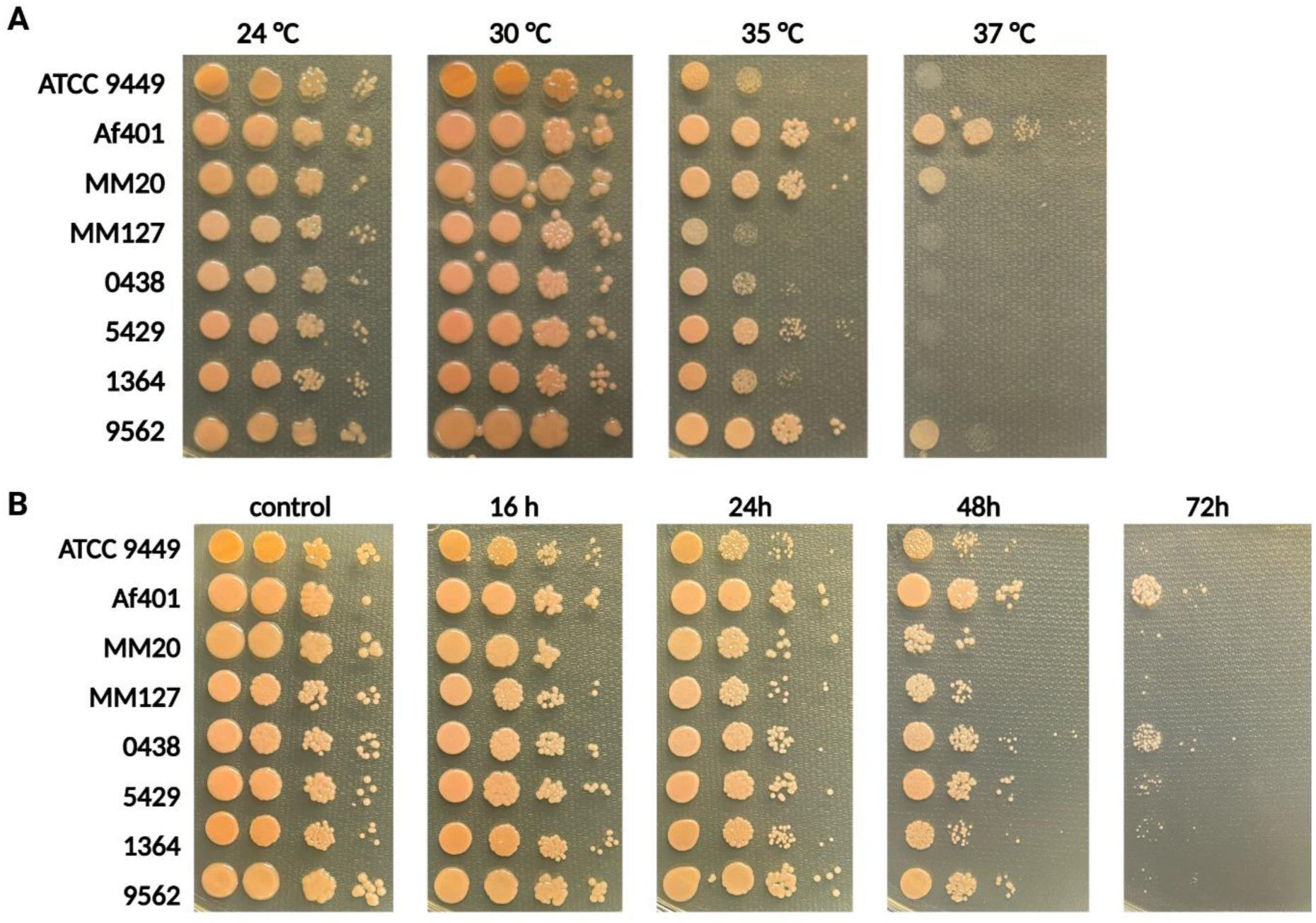
Temperature-dependent growth and thermotolerance of *Rhodotorula mucilaginosa* strains. (A) Growth of selected *R. mucilaginosa* strains at different temperatures. Tenfold serial dilutions of yeast cells were spotted onto YPD agar and incubated at 24, 30, 35, or 37°C for 48 h to assess growth under environmental, optimal, and host-associated temperature conditions. (B) Thermotolerance assay of the same *R. mucilaginosa* strains at 42°C. Tenfold serial dilutions of yeast cells were spotted onto YPD agar and incubated at 42°C for 16, 24, 48, or 72 h. Plates were subsequently transferred to 30°C and incubated for 48 h to allow growth of surviving cells. Control plates were maintained at 30°C throughout the experiment. Growth following recovery was used as a measure of survival after exposure to elevated temperature.

All strains presented very similar growth on YPD agar media when incubated at 24°C and 30°C, indicating no visible differences in viability under environmental or optimal laboratory growth conditions (Fig. 1A). However, growth differences became apparent at 35°C. Laboratory strain ATCC 9449, marine mammal isolate MM127, and clinical strains 0438 and 1364 displayed a noticeable reduction in growth relative to that observed at lower temperatures, whereas mosquito strain Af401, marine mammal strain MM20, and clinical strains 5429 and 9562 showed little or no growth impairment under the same conditions (Fig. 1A). Incubation at 37°C resulted in a marked reduction in growth for nearly all strains tested, with the exception of strain Af401 which showed only minor reduction in growth at 37°C compared to lower temperatures, suggesting that this strain has a greater capacity to tolerate host body temperature compared to the other tested strains of *R. mucilaginosa* (Fig. 1A).

We next evaluated the ability of *R. mucilaginosa* strains to withstand and recover from heat stress after exposure to temperatures associated with febrile conditions. Cells were spotted onto YPD agar plates, exposed to 42°C for 16-72 h, and subsequently allowed to recover to 30°C for 48 hr (Fig. 1B). Exposure to 42°C for 16 or 24 h had only a limited effect on the viability of the tested strains, as evidenced by robust growth following subsequent incubation at 30°C (Fig. 1B). After 48 h of incubation at 42°C, most strains displayed reduced growth, indicating decreased survival after prolonged heat stress. The greatest impact was observed after 72 h of exposure to 42°C. Under these prolonged high-temperature stress conditions, nearly all strains halted growth and were unable to recover, demonstrating a severe impact of heat on cell viability. Interestingly, strains Af401 and 0438 retained some limited growth after 72 h of heat stress, indicating improved heat tolerance in these isolates (Fig. 1B).

Taken together, we identified strain-specific differences in temperature tolerance and ability to recover following heat stress, with no clear trends between clinical vs. non-clinical isolates. All tested *R. mucilaginosa* strains tolerated short-term exposure to temperatures associated with extreme fever, but prolonged exposure to 42°C severely affected the viability of all isolates. Strain Af401, isolated from a mosquito in Zambia, exhibited the highest degree of thermotolerance, with less growth impairment at 37°C compared to other strains, and displayed limited ability to recover growth after prolonged heat stress. Clinical isolate 0438 exhibited growth impairment when incubated at 35° and 37°C for 48 h, yet retained some ability to recover after heat stress at 42°C once returned to more favorable conditions.

### Temperature differentially affects biofilm biomass and metabolic activity in *Rhodotorula mucilaginosa*

In the next phase of this study, we analyzed the impact of temperature on static biofilm formation using modifications of previously published protocols. Selected *R. mucilaginosa* strains were incubated in 96-well polystyrene plates in RPMI 1640 medium for 3 days at 24°, 30°, or 37°C, and biofilm development biomass was quantified by crystal violet (CV) staining, and biofilm metabolic activity was quantified via XTT reduction assays (Fig. 2).

**Figure 2.**
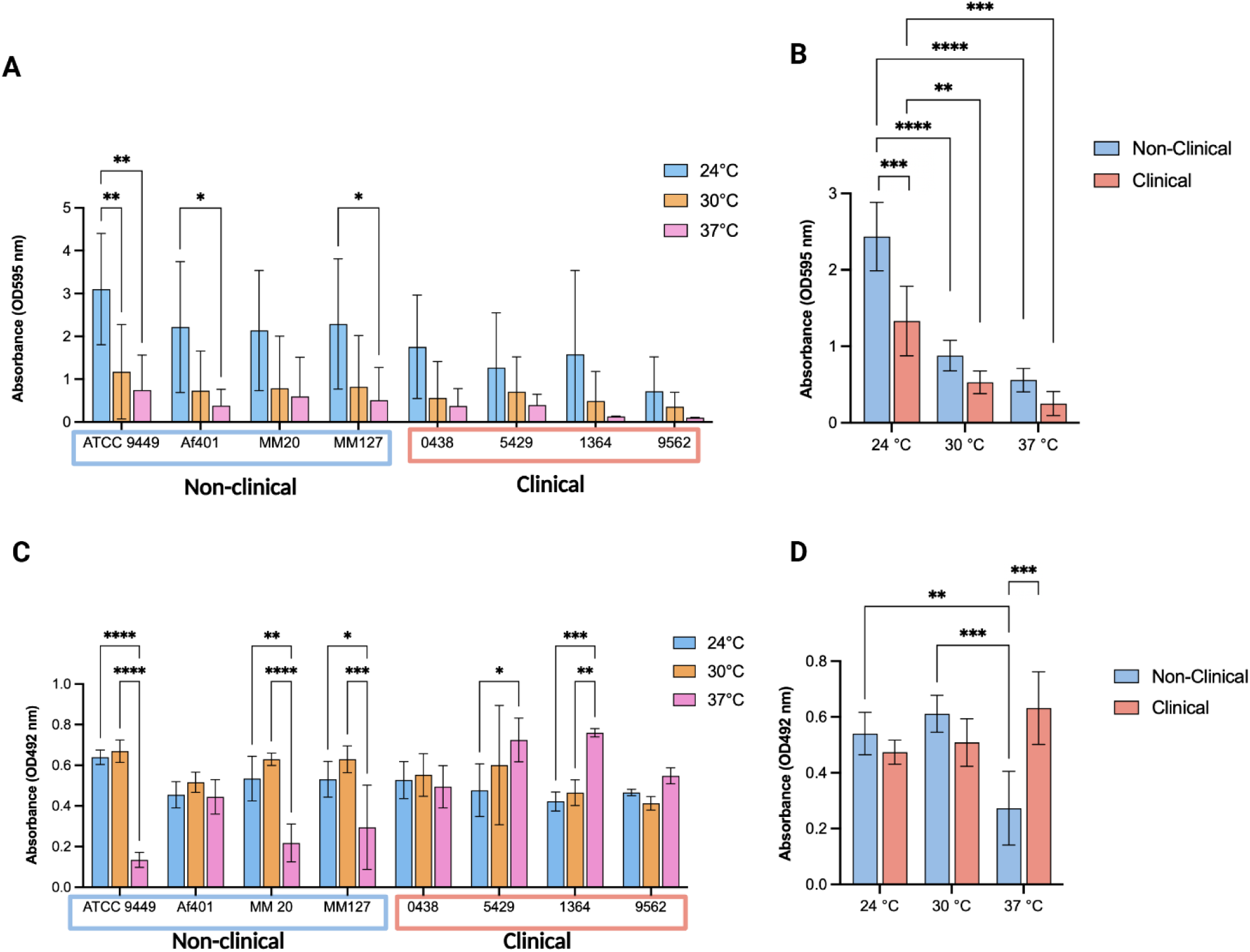
Temperature-dependent biofilm formation by *Rhodotorula mucilaginosa* strains. Selected *R. mucilaginosa* strains were cultured in RPMI 1640 medium in 96-well plates for 3 days at 24°C, 30°C, or 37°C. (A) Biofilm biomass was quantified by crystal violet staining in the form of a bar graph. (B) Bar graph presenting the difference in biofilm mass quantification for combined Clinical and Non-clinical strains. (C) Biofilm metabolic activity was assessed using the XTT reduction assay. (D) Bar graph presenting the difference in biofilm metabolic activity for combined Clinical and Non-clinical strains. Data represent means ± standard deviations. Statistical significance was determined by two-way ANOVA. Significance is denoted as *p < 0.05, **p < 0.01, ***p < 0.001, ****p < 0.0001

Several tested strains (ATCC 9449, Af401, MM127) produced biofilm biomass more efficiently at room temperature compared to higher temperatures, as assessed by CV staining (Fig. 2A). When comparing pooled data for clinical vs. non-clinical isolates, non-clinical strains had larger biofilm biomass at 24 °C compared to clinical isolates, while no differences in isolate type were apparent at higher temperatures (Fig. 2B). Additionally in the pooled results we observed that that for clinical and non-clinical isolates, biofilm biomass at 24°C was greater than biofilm biomass at 30°C or 37°C (Fig. 2B)

Analysis of biofilm metabolic activity by XTT reduction assay presented a more complex, strain-dependent response to various incubation temperatures (Fig. 2C). Laboratory strains ATCC 9449 and marine mammal strains MM20 and MM127 showed significantly higher metabolic activity in biofilms formed at lower temperatures (24 and 30°C) than in biofilms formed at high temperature (37°C). In contrast, this pattern was not observed for clinical isolates. Clinical strains 0438 and 9562 showed no significant differences in biofilm metabolic activity between the tested temperatures, whereas strains 5429 and 1364 produced biofilms with significantly higher metabolic activity at 37°C compared to 24°C. When pooled data for non-clinical and clinical isolates were compared, no differences in metabolic activity were observed between biofilms formed at 24 °C or 30 °C, whereas at 37 °C, biofilms from clinical isolates showed significantly greater metabolic activity than biofilms from non-clinical isolates (Fig 2D). These results suggest that clinical isolates can maintain biofilm biomass and sustain biofilm activity across a wider range of temperatures, whereas mammalian body temperature inhibited biofilm formation in the laboratory strain ATCC 9449 and the two isolates from harbor seal nasal cavities. Interestingly, the mosquito isolate Af401 showed a similar pattern to the clinical isolates with no differences in biofilm formation between temperatures.

### Cell surface hydrophobicity

Cell-surface hydrophobicity may be important in initial surface attachment and promotion of biofilm development, as observed in other fungal species like *Candida spp.* or *Cryptococcus spp*. (29–31). We evaluated the surface hydrophobicity of selected *R. mucilaginosa* strains using a microbial adhesion to hydrocarbons (MATH) assay, which indicates the proportion of yeast cells that adhere to the non-polar solvent. All strains exhibited moderate to high levels of hydrophobicity, although the degree of hydrophobicity varied among isolates (Fig. 3).

**Figure 3.**
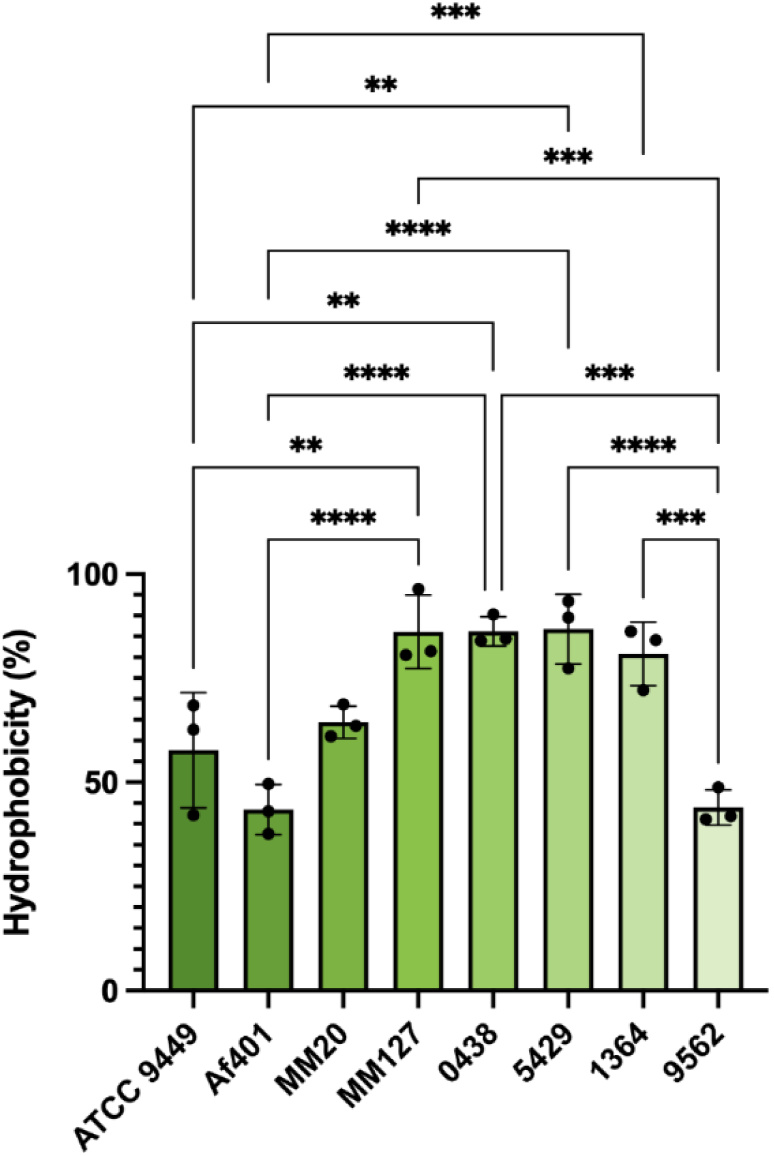
Cell surface hydrophobicity. Cell surface hydrophobicity of the selected *R. mucilaginosa* strains. Hydrophobicity was determined using the microbial adhesion to hydrocarbons (MATH) assay. Data are presented as means ± standard deviations from independent experiments. Significance is denoted as *p < 0.05, **p < 0.01, ***p < 0.001, ****p < 0.0001

### Biofilm formation on polyurethane intravenous catheters

Next, we evaluated the efficiency of biofilm formation of various *R. mucilaginosa* strains on the surface of sterile polyurethane IV catheters (Fig. 4). Catheter fragments (10mm) were incubated with fungal cultures for 3 days at 30°C under dynamic conditions, washed to remove nonadherent cells, and analyzed using the XTT reduction assay to quantify metabolically active cells. The results revealed substantial variation in initial biofilm formation among tested strains. Strains Af401, 5429, and 1364 produced the highest XTT signals, relative to the laboratory strain ATCC 9449, suggesting that these strains either have better capacity to colonize polyurethane catheter material or that the biofilms present are more metabolically active (Fig. 5A).

**Figure 4.**
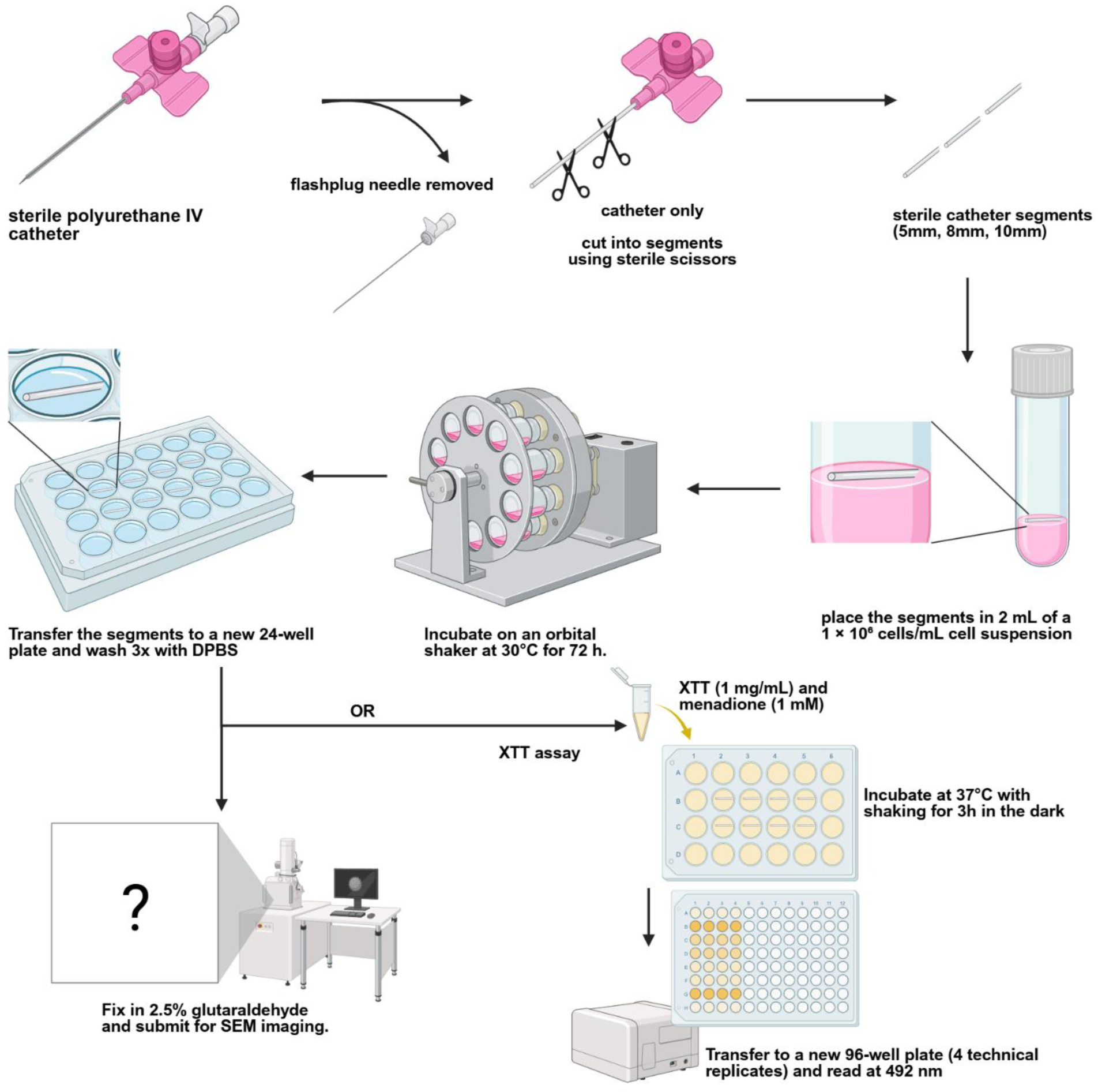
Experimental workflow for biofilm formation on polyurethane intravenous (IV) catheters. Schematic illustration of the experimental procedure used to induce *R. mucilaginosa* biofilm formation on polyurethane IV catheters. Catheter segments were cut into 10-mm fragments, incubated with *R. mucilaginosa* cultures under agitation for 3 days at 30°C to promote biofilm formation, washed to remove nonadherent cells, and subsequently processed for scanning electron microscopy (SEM) or XTT reduction assays to evaluate biofilm formation.

**Figure 5.**
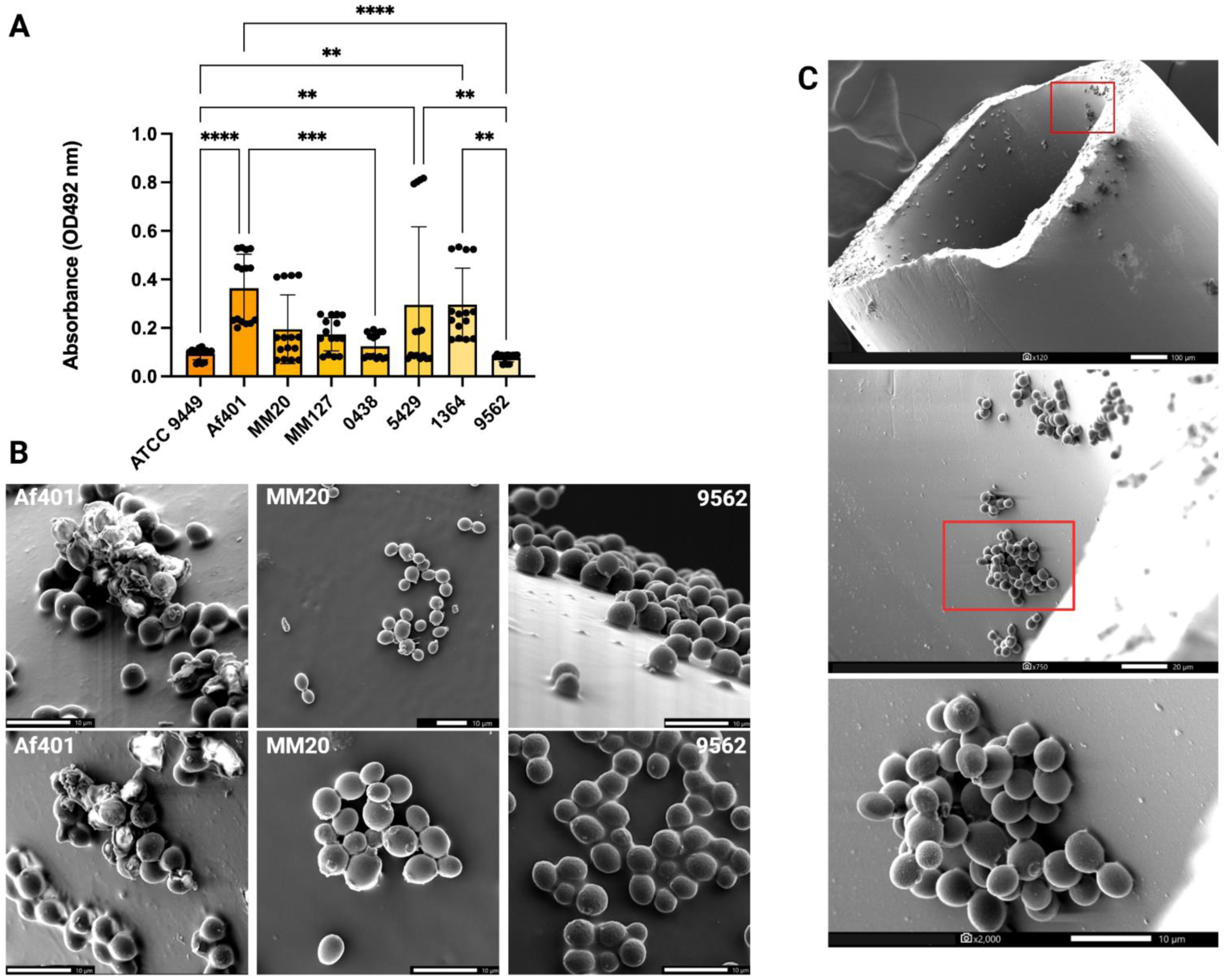
Biofilm formation by *R. mucilaginosa* on polyurethane intravenous (IV) catheters. (A) Quantification of biofilm formation by selected *R. mucilaginosa* strains on polyurethane IV catheters. Catheters were incubated in agitated cell cultures for 3 d at 30°C, and biofilm metabolic activity was subsequently assessed using the XTT reduction assay. Data are presented as means ± standard deviations from independent experiments. (B) Representative scanning electron microscopy (SEM) images of *R. mucilaginosa* biofilms formed on polyurethane IV catheter surfaces during the interfacial aggregation stage of biofilm development. Images are shown for strains representing different origins: clinical isolate 9562, marine mammal isolates MM20, and environmental isolate AF401. Scale bars, 10 μm. (C) Representative SEM image (120×, 750×, and 2000× magnification) of the inner surface of a polyurethane IV catheter colonized by *R. mucilaginosa*. Significance is denoted as *p < 0.05, **p < 0.01, ***p < 0.001, ****p < 0.0001

Following the biochemical evaluation of biofilm formation on polyurethane IV catheters, we visualized the results for three selected strains by SEM: environmental isolates Af401 and MM20, and clinical isolate 9562 (Fig. 5B). SEM analysis confirmed the presence of adherent *R. mucilaginosa* cells on the catheter surface and revealed the formation of multicellular aggregates characteristic of the interfacial aggregation stage of biofilm development. These observations support the XTT-based findings and demonstrate that *R. mucilaginosa* strains can adhere to and colonize polyurethane IV catheter material, although the extent of biofilm formation varies among isolates. Further, we investigated the ability of *R. mucilaginosa* to adhere to and initiate biofilm formation on the inner catheter surface. Adherent cells were frequently observed in multicellular aggregates rather than as isolated yeast cells. Higher-magnification imaging demonstrated densely packed clusters of yeast cells attached to the catheter surface, consistent with the interfacial aggregation stage of biofilm development (Fig. 5C). These observations further confirm that *R. mucilaginosa* readily adheres to polyurethane catheter material and can initiate biofilm formation on the surfaces of clinically relevant plastic medical devices.

### Pigmentation analysis of *R. mucilaginosa*

Given the role of carotenoids in conferring UV-C resistance, we compared pigment intensity among the tested *R. mucilaginosa* strains by quantifying colony color after 48 h of growth on YPD media using the CIELAB color space (Fig. 6). Analysis of colony lightness (L*) revealed comparable pigmentation intensity across all strains. Similarly, only minor differences were observed in the red–green coordinate (a*). In contrast, analysis of the yellow–blue coordinate (b*) identified strain ATCC 9449 as the only isolate exhibiting significantly greater yellow pigmentation than the remaining strains. These results confirm our initial visual assessment that colony pigmentation is largely comparable among the tested *R. mucilaginosa* isolates.

**Figure 6.**
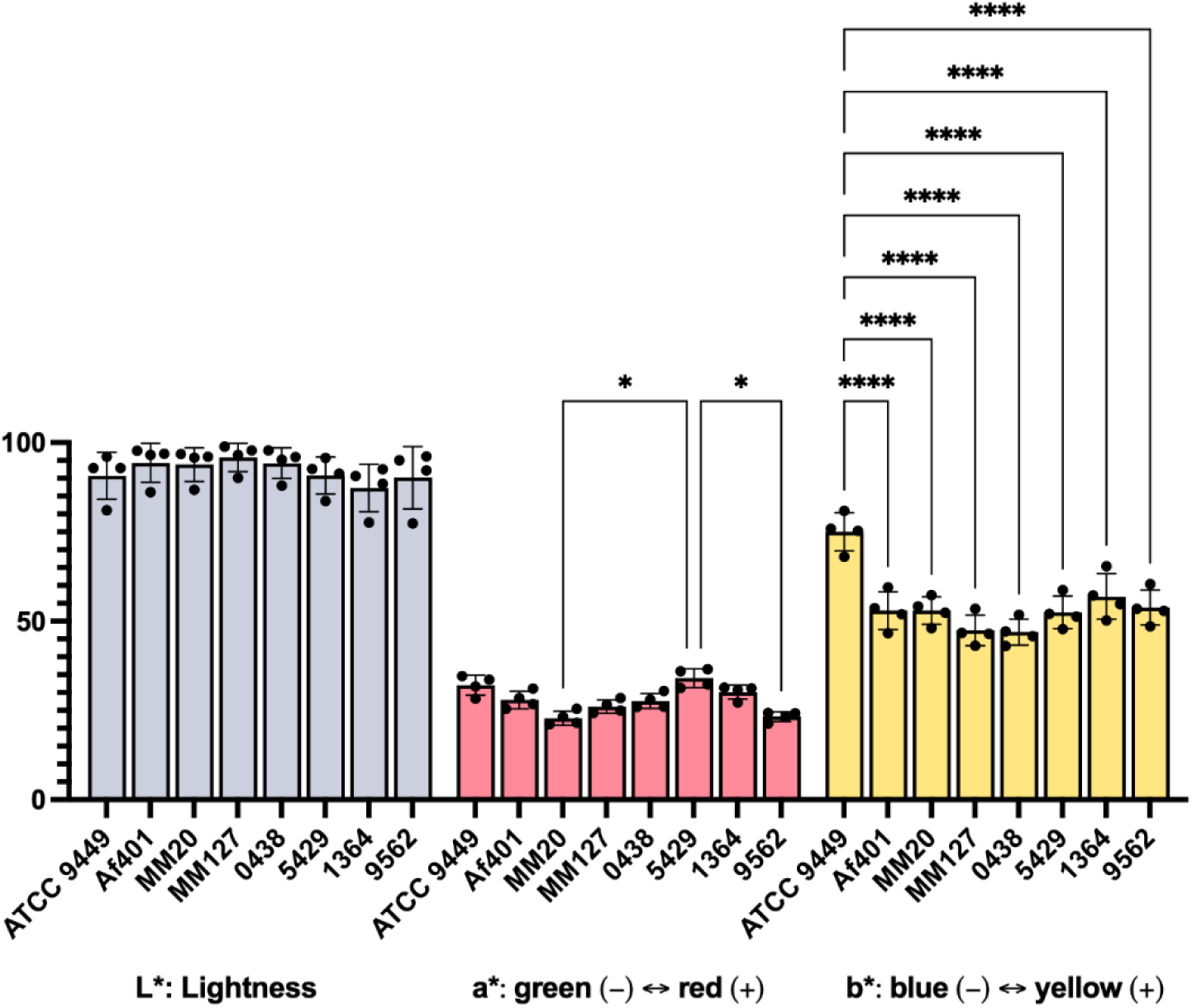
Quantitative analysis of colony color. Colony color was quantified from images of colonies after 48 h of incubation using the CIELAB color space. Lightness (L*). Red–green coordinate (a*), and Yellow–blue coordinate (b*). Bars represent the mean ± SD of four independent experiments (n = 4). Statistical analysis was performed using two-way ANOVA. Significance is denoted as *p < 0.05, ****p < 0.0001

### Biofilms enhance tolerance of *R. mucilaginosa* to UV-C exposure

Ultraviolet (UV) radiation is commonly used for surface sterilization in clinical and research settings. Biofilms have been shown to confer UV-C resistance in other microbial species (32–35). To determine the baseline susceptibility of *R. mucilaginosa* to UV exposure, planktonic cells from eight strains were subjected to increasing doses of UV radiation, and survival was evaluated by spot dilution assay (Fig. 7A). Growth progressively decreased with increasing UV-C dose in all tested Rhodotorula strains. Exposure to 50 mJ/cm^2^ resulted in a marked reduction in growth in all strains, consistent with substantially impaired planktonic cell viability. A similar evaluation of UV susceptibility for *C. albicans*, *C. neoformans* and *C. gattii* confirmed previous observations that those opportunistic fungal pathogens are much more susceptible to UV-C irradiation, presenting dramatic growth inhibition at 30 mJ/cm^2^ for *C. albicans* and 40 mJ/cm^2^ for *C. neoformans* and *C. gattii* (Fig 7B).

**Figure 7.**
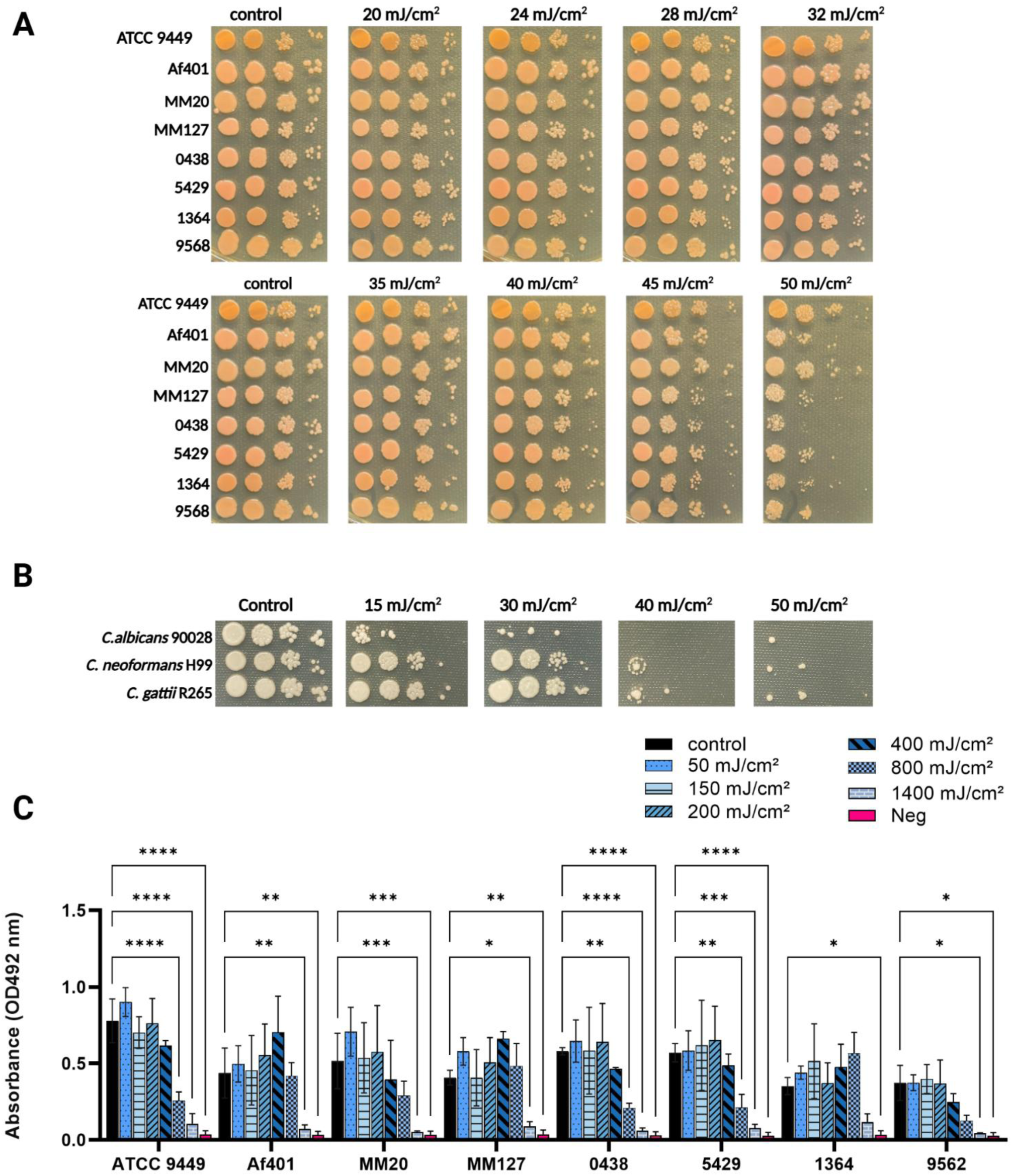
Analysis of UV-C resistance in planktonic and biofilm yeast cells. (A) UV-C susceptibility of Rhodotorula planktonic cells. Tenfold serial dilutions of selected *R. mucilaginosa* strains were spotted onto YPD agar and exposed to increasing doses of UV radiation (0, 20, 24, 28, 32, 35, 40, 45, and 50 mJ/cm^2^). Following UV exposure, plates were incubated at 30°C to assess cell survival and growth. UV-C susceptibility of representative strains of selected yeasts, *Candida albicans*, *Cryptococcus neoformans*, and *Cryptococcus gattii*, to increasing doses of UV radiation (0, 15, 20, 30, 40, and 50 mJ/cm^2^) was assessed using a tenfold serial dilution spot assay. (C) UV-C susceptibility of biofilm-associated cells. Selected *R. mucilaginosa* strains were grown as biofilms in RPMI 1640 medium in static 96-well plates for 3 days at 30°C. Mature biofilms were exposed to UV radiation at doses of 0, 50, 150, 200, 400, 800, or 1400 mJ/cm^2^ and subsequently incubated in fresh RPMI 1640 medium for 24 h at 30°C. Negative control illustrates results for biofilm with cells inactivated by 20 h incubation in 70% ethanol. Biofilm viability was then determined using the XTT reduction assay. Data are presented as means ± standard deviations from independent experiments. Statistical significance was determined by two-way ANOVA. Significance is denoted as *p < 0.05, **p < 0.01, ***p < 0.001, ****p < 0.0001

We next examined whether the UV dose effective against *R. mucilaginosa* planktonic cells was sufficient to reduce the viability of biofilm-associated cells. Mature biofilms were exposed to UV doses ranging from 50 to 1400 mJ/cm^2^, and viability was subsequently assessed using the XTT reduction assay (Fig. 7C). In contrast to planktonic cells, biofilm-associated cells remained largely unaffected by UV exposure. No significant reduction in biofilm viability was observed after treatment with 50, 150, 200, or 400 mJ/cm^2^ of UV-C radiation compared with untreated controls. In strains ATCC 9449, 0438, and 5429, a dosage of 800 mJ/cm^2^ was necessary to observe a statistically significant reduction of biofilm viability, while in strains Af401, MM20, MM127, and 9562, a dosage of 1400 mJ/cm^2^ was necessary to achieve a similar effect. Strain 1364 presented a visible but not statistically significant reduction in biofilm metabolic activity even at the highest level of UV-C exposure. These results reveal a marked difference in susceptibility to UV-C between planktonic and biofilm-associated *R. mucilaginosa* cells, with biofilm growth associated with increased resistance to UV-mediated killing.

## Discussion

High body temperature is widely recognized as one of the barriers preventing environmental fungi with pathogenic potential from causing disease in mammals (36, 37). Increasing reports of *R*. *mucilaginosa* infections raise the possibility that enhanced thermotolerance could contribute to the emergence of strains with greater pathogenic potential. Climate warming has been proposed to select for environmental fungi capable of growth at progressively higher temperatures, thereby narrowing the thermal barrier that protects mammals from fungal disease (37–39). Since *R. mucilaginosa* has emerged as an increasingly important opportunistic fungal pathogen, relatively little is known about the thermal constraints that limit its growth within the host. In the present study, all isolates grew optimally at 30°C, and most exhibited reduced growth at 35°C, indicating that temperatures encountered at peripheral body sites already impose a constraint on fungal viability. Despite their limited capacity to proliferate at 37°C, all isolates displayed considerable tolerance to acute heat stress. Fever may function as a host defense mechanism by creating a thermal barrier that severely inhibits the growth and survival of thermally sensitive fungal pathogens (40). Exposure to 42°C for up to 48 h produced only a modest reduction in viability, and no consistent differences were observed between environmental and clinical isolates. Only prolonged exposure for 72 h resulted in near-complete loss of viability in most strains, suggesting that *R. mucilaginosa* can withstand limited exposure to extreme febrile-like temperatures despite its restricted ability to sustain growth at host body temperature. This illustrates its noteworthy pathogenic potential as an emerging fungal pathogen and threat to public health.

Temperature also influenced biofilm development. Quantification of total biofilm biomass by crystal violet staining showed a potential trend across the strain collection, with biofilms formed at 24°C generally exhibiting the greatest biomass, while biofilms formed at 37°C accumulated less biomass. This observation suggests that environmental temperatures favor biofilm production, which may facilitate persistence outside the host. In contrast, assessment of biofilm metabolic activity revealed a different pattern. Three of the four nonclinical isolates exhibited their lowest metabolic activity in biofilms formed at 37°C, consistent with the overall reduction in fungal growth observed at host temperature. In comparison, two of the four clinical isolates maintained significantly greater biofilm metabolic activity at 37°C than at lower temperatures. Although the number of isolates examined is limited, this finding raises the possibility that clinical isolates have undergone selection for improved metabolic performance under host-associated conditions. Host-adaptation could permit survival and persistence of fungi even when overall fungal proliferation is constrained by mammalian body temperature. While the thermal barrier is a major determinant limiting *R. mucilaginosa* pathogenicity, our results support the hypothesis that adaptation to host temperature may occur in clinical strains, enhancing fungal thermotolerance and improving cell survival.

Biofilm formation on indwelling medical devices is a defining feature of many opportunistic fungal pathogens and is thought to contribute substantially to the increasing incidence of *Rhodotorula* catheter-associated bloodstream infections (9, 13, 25, 41). Although *R. mucilaginosa* has frequently been isolated from contaminated central venous catheters, quantitative methods for assessing its ability to colonize clinically relevant biomaterials remain limited (6, 15). Consequently, we sought to characterize biofilm formation on polyurethane intravenous catheters and to establish a simple, reproducible assay to compare the biofilm-forming capacity of different *R. mucilaginosa* isolates. Our initial evaluation with scanning electron microscopy provided direct evidence that *R. mucilaginosa* readily adheres to the inner surface of polyurethane catheters and forms multicellular aggregates characteristic of the early stages of biofilm development. These observations are consistent with previous reports implicating catheter colonization as a critical step in the pathogenesis of *Rhodotorula* infections. In parallel, analysis of cell surface hydrophobicity revealed that most isolates exhibited moderate to high hydrophobicity, a property that is frequently associated with enhanced adhesion to abiotic surfaces. Although cell surface hydrophobicity alone is unlikely to determine biofilm formation, these findings suggest that the surface characteristics of *R. mucilaginosa* are compatible with efficient attachment to catheter materials. To complement the qualitative SEM observations, we modified an XTT-based assay to quantify metabolically active biofilms formed directly on polyurethane catheter fragments and confirmed that the tested strains can form biofilm complexes on this material. Furthermore, low variability of the results among biological replicates indicates good reproducibility of this assay for further evaluation of clinical isolates. Importantly, Af401, the *R. mucilaginosa* strain that produced the highest XTT signals also exhibited more robust biofilm formation based on SEM evaluation. Because the assay is simple, reproducible, and compatible with standard laboratory equipment, it may serve as a useful platform for future studies to compare the biofilm-forming capacity of clinical isolates, identify strains with enhanced device-colonizing potential, and evaluate strategies to prevent catheter-associated *Rhodotorula* infections.

Ultraviolet irradiation is a widely used method for decontamination of surfaces in healthcare facilities and during the manufacturing of biomedical devices because of its effectiveness against a broad range of microorganisms (42–47). Production of carotenoids is one of the strategies utilized by microorganisms to enhance protection against UV-C radiation (24, 48, 49). The evaluation of our tested strains for potential differences in colony pigmentation indicated rather comparable intensity of pigmentation across all tested strains. All tested *R. mucilagionasa* strains exhibited substantial resistance to UV exposure, with marked growth inhibition observed only at the highest doses tested. This level of UV-C (254nm) far exceeds the dosage required for inactivation of planktonic bacterial species *Staphylococcus aureus* or *Pseudomonas aeruginosa* (5mJ/cm2) or fungal species *Candida albicans* and *Candida parapsilosis* (40 mJ/cm^2^), as well as previously reported for non-melanized strains of *Cryptococcus neoformans* and *Cryptococcus gattii* (50–53). The observed UV tolerance is consistent with previous studies demonstrating that carotenoid pigments produced by *R. mucilaginosa* may protect cells from UV-induced damage (48, 49). We then asked whether biofilm formation provides additional protection against UV exposure. Although a UV dose of 50 mJ/cm^2^ markedly reduced the viability of planktonic cells, biofilm-associated cells remained largely unaffected, not only at this dose but also following exposure to doses as high as 400 or 800 mJ/cm². Thus, biofilm growth conferred protection against UV-C doses approximately fourfold greater than those sufficient to impair planktonic cells. These findings confirmed similar observations in other fungal species that the extracellular biofilm environment provides additional protection beyond the intrinsic UV resistance of individual *Rhodotorula* cells (54). The mechanisms responsible for this enhanced resistance were not investigated in the present study but may include attenuation of UV-C penetration by the biofilm matrix together with physiological adaptations associated with biofilm growth. Similar protective effects of fungal biofilms against antimicrobial agents and environmental stresses have been described for other opportunistic fungal pathogens and could be studied in *R. mucilaginosa* in future work (32, 33, 35, 55, 56).

Collectively, our findings identify biofilm formation as an important determinant of *R. mucilaginosa* environmental persistence. In addition to facilitating colonization of indwelling medical devices, biofilms substantially reduce the susceptibility of *R. mucilaginosa* to UV-mediated sterilization. These observations have practical implications for infection control, as they suggest that standard UV decontamination protocols effective against planktonic cells may be insufficient to eliminate *Rhodotorula* biofilms from contaminated medical devices or manufacturing environments.

## Materials and Methods

### Fungal strains and culture conditions

*Rhodotorula mucilaginosa* ATCC 9449 and Af401 were obtained from Daniel F. Q. Smith. *R. mucilaginosa* strains 0438, 5429, 1364, and 9562 were isolated from blood cultures of hospitalized patients as part of clinical management and obtained from Sean Zhang at the Johns Hopkins Hospital. *R. mucilaginosa* strains MM20 and MM127 were isolated by IAJ from nasal swabs collected from harbor seals in collaboration with staff at Sealife Response, Rehabilitation, and Research (SR3) under NOAA permit #24359. *Candida albicans* 90028, *Cryptococcus neoformans* H99, and *Cryptococcus gattii* R265 from Casadevall laboratory stocks. All strains used in this study were maintained as glycerol stocks at − 80°C. For routine culture, strains were streaked onto YPD (BD Difco™, Becton, Dickinson and Company, Sparks, MD, USA) agar plates from frozen stocks and incubated overnight at 30°C. Plates were sealed with Parafilm and stored at 4°C until use. For experimental use, cells were inoculated into 2 mL YPD liquid medium and cultured overnight at 30°C on an orbital shaker. We used overnight cultures for all subsequent experiments (unless otherwise specified).

### Spot Assays for Growth and Stress Tolerance

Overnight YPD *R. mucilaginosa* cultures were pelleted by centrifugation at 4000 rpm for 4 min, washed once with sterile Dulbecco’s phosphate-buffered saline (DPBS; Corning, Manassas, VA, USA), and resuspended in DPBS. Cell suspensions were adjusted to an approximate concentration of 3.3 × 10^6^ cells/mL and subjected to 10-fold serial dilutions (10^0^-10^−3^) in DPBS using a sterile 96-well plate (CELLTREAT, Pepperell, MA, USA). A volume of 3 μL of each dilution was spotted onto YPD plates using a multichannel pipette and allowed to air-dry before incubation. Plates were incubated under the indicated conditions and imaged after 2–3 days.

For thermotolerance analysis, spotted plates were incubated at 24, 30, 35, or 37°C for 48 h and subsequently photographed to assess growth under different temperature conditions. To analyze resistance to heat stress, spotted plates were incubated at 42°C for 16, 24, 48, or 72 h and subsequently transferred to 30°C for approximately 48 h of recovery before imaging.

For measurement of UV-C tolerance, spotted plates were exposed to UV-C irradiation (254 nm) at doses of 20, 24, 28, 32, 35, 40, 45, or 50 mJ/cm² using a Stratalinker 1800 UV crosslinker (Stratagene, La Jolla, CA) with the plate lids removed. Following irradiation, plates were transferred to 30°C and incubated for approximately 48 h to allow recovery before imaging.

### Microbial Adhesion to Hydrocarbons (MATH) Assay (Cell Surface Hydrophobicity Assay)

The evaluation of the cell-surface hydrophobicity of *R. mucilaginosa* strains was performed according to published protocols (57–59). Cell cultures were grown overnight in 5 mL YPD broth at 30°C. Cells were harvested by centrifugation (2,500 xg, 5 min), washed three times with DPBS, and resuspended in at least 3.5 mL DPBS. Cell suspensions were adjusted to an OD600 of 0.3–0.4. Aliquots (100 μL) were transferred to a 96-well plate in triplicate, and OD600 was measured using a SpectraMax iD5 microplate reader (Molecular Devices, San Jose, CA, USA) to obtain the initial absorbance value (A0). Subsequently, 3 mL of the cell suspension was transferred to a glass tube and 500 μL n-hexadecane (Sigma-Aldrich, St. Louis, MO, USA) was added. Tubes were sealed with Parafilm and vortexed for 60 s before being incubated at room temperature for 2 min to allow phase separation. Following separation, 100 μL of the lower aqueous phase was transferred into a 96-well plate in triplicate, and OD600 was measured to obtain A1.

Cell surface hydrophobicity (%CSH) was calculated as:

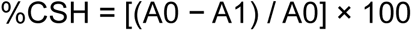

### Static biofilm formation

Biofilm production was adapted from the previously described method. (57, 60) Cells were collected and washed as described above, and resuspended in RPMI 1640 medium (Gibco, Thermo Fisher Scientific, Waltham, MA, USA). Cell suspensions were prepared by two sequential 1:10 dilutions for cell counting and then adjusted to a final concentration of 1 × 10^6^ cells/mL in RPMI 1640 medium. Biofilm formation was tested in sterile 96-well polystyrene plates. A total of 200 μL of the adjusted cell suspension was added to each well, followed by incubation at 24°C, 30°C, or 37°C for 72 h. For each strain and condition, four technical replicates were included in each experiment. Experiments were independently repeated three to five times.

### Biofilm biomass quantification

Crystal violet staining was adapted and modified based on the published protocols (57, 61). Following biofilm formation, wells were gently washed twice with 200 μL DPBS to remove non-adherent cells. Biofilms were stained with 110 μL of 0.4% (w/v) crystal violet solution prepared in DPBS from a stock solution (Sigma-Aldrich, St. Louis, MO, USA) for 45 min. Following staining, each well was washed four times with 200 μL DPBS to remove excess dye. The stained biofilms were then destained with 200 μL of 95% ethanol for 45 min. Subsequently, 100 μL of the destaining solution from each well was transferred to fresh wells, and absorbance was measured at 595 nm using a microplate reader. Each strain was tested independently four times on different days.

### Biofilm metabolic activity

XTT reduction assay was adapted and modified based on the published protocols (54, 62). A semiquantitative measurement of *R. mucilaginosa* biofilm formation was obtained from the 2,3-bis(2-methoxy-4-nitro-5-sulfophenyl)-5-[(phenylamino) carbonyl]-2H-tetrazolium-hydroxide (XTT) reduction assay. For *R. mucilaginosa* strains, 100 μl of XTT salt solution (1 mg/mL in Milli-Q water; Cayman Chemical, Ann Arbor, MI, USA, or Invitrogen Molecular Probes, Eugene, OR, USA) and 8 μl of menadione solution (1 mM in acetone; Sigma-Aldrich, St. Louis, MO, USA) were added to each well. Following incubation, 80 μL aliquots were transferred from each well to fresh wells. Absorbance was then measured at 492 nm using a microplate reader.

### UV Tolerance of Established Static Biofilms

Static biofilms used for UV tolerance assays were established at 24°C as described above. Following biofilm formation, wells were washed twice with DPBS and allowed to air dry for 5 min. Biofilms were exposed to UV-C radiation (254 nm) at doses of 50, 150, 200, 400, 800, and 1400 mJ/cm^2^ with the plate lid removed. Following irradiation, 200 μL fresh RPMI 1640 medium was added to each well, and biofilms were allowed to recover at 24°C for 20 h without shaking. The negative control was performed for biofilm with cells inactivated by 20 h incubation in 70% ethanol. After recovery, wells were washed twice with DPBS, and biofilm metabolic activity was quantified using the XTT reduction assay as described above.

### CIELAB color analysis

Colony color was analyzed using the CIELAB color space. Spot assay plates were incubated at 30°C for 48 h before imaging. All photographs were captured under standardized conditions using the same smartphone. Images in HEIC format were converted to TIFF format and standardized to the RGB color space prior to analysis. The TIFF images were then analyzed in Fiji (ImageJ, version 1.54p) using the RGB to CIELAB plugin, which converted the images from RGB to CIELAB and generated L*, a*, and b* values. For each image, an elliptical region of interest (ROI) was manually drawn over the largest spot. Regions containing visible specular reflections were excluded from the ROI. Mean L*, a*, and b* values were recorded for each ROI. Four independent biological experiments were analyzed, with one ROI measured per image.

### Dynamic biofilm formation on intravenous catheter

*R. mucilaginosa* biofilms were grown on sterile polyurethane IV catheter (20G × 1; Covetrus, Dublin, OH, USA) segments under dynamic conditions. Sterile catheter segments (1 cm in length) were placed into capped culture tubes containing 2 mL RPMI 1640 medium inoculated with *R. mucilaginosa* at a final concentration of 1 × 10^6^ cells/mL. The cultures were incubated at 30°C on an orbital shaker for 72 h to allow biofilm formation on the catheter surfaces.

Following incubation, the catheter segments were transferred to a sterile 24-well plate and washed three times with DPBS to remove non-adherent cells. Catheter-associated biofilms were quantified using the XTT reduction assay as described above. Following washing, catheter segments were incubated with 500 μL XTT solution and 40 μL menadione solution at 37°C for 3 h in the dark. Following incubation, 100 μL of the reaction mixture was transferred into four wells of a new 96-well plate as technical replicates, and absorbance was measured at 492 nm using a microplate reader.

### Scanning Electron Microscopy (SEM)

For SEM imaging, biofilms were grown on 5 mm and 8 mm IV catheter segments. Samples were fixed in 2.5% glutaraldehyde, 3mM MgCl2, in 0.1 M sodium cacodylate buffer, pH 7.2 overnight at 4C. After buffer rinse, samples were postfixed in 1% osmium tetroxide in 0.1 M sodium cacodylate buffer for 30 m at room temperature, protected from light. Following a DH2O rinse, samples were dehydrated in a graded series of ethanol and left to dry overnight with hexamethyldisilazane (HMDS). Samples were mounted on carbon-coated stubs, coated with 5nm gold palladium (Denton DeskV), and imaged on a JEOL SEM (JSM-IT700HR) at 5 kV.

## Acknowledgments

We thank Daniel F. Q. Smith and Sean Zhang for providing access to selected strains for this study and for advising us on methodology. We thank Barbara Smith at the Microscope Facility at the Johns Hopkins University School of Medicine for performing the SEM imaging.

A.C. was supported in part by National Institutes of Health R01 Grants HL059842, AI152078, AI052733, and U19AI189183.

